# Single-vessel cerebral blood flow fMRI to map blood velocity by phase-contrast imaging

**DOI:** 10.1101/2020.09.03.280636

**Authors:** Xuming Chen, Yuanyuan Jiang, Sangcheon Choi, Rolf Pohmann, Klaus Scheffler, David Kleinfeld, Xin Yu

**Author notes:** Lead Contact: Address: Max-Planck-Ring. 11, 72076, Tübingen, Germany (Xin Yu).

## Abstract

Current approaches to high-field fMRI provide two means to map hemodynamics at the level of single vessels in the brain. One is through changes in deoxyhemoglobin in venules, i.e., blood oxygenation level-dependent (BOLD) fMRI, while the second is through changes in arteriole diameter, i.e., cerebral blood volume (CBV) fMRI. Here we introduce cerebral blood flow (CBF)-fMRI, which uses high-resolution phase-contrast MRI to form velocity measurements of flow and demonstrate CBF-fMRI in single penetrating microvessels across rat parietal cortex. In contrast to the venule-dominated BOLD and arteriole-dominated CBV fMRI signal, the phase-contrast -based CBF signal changes are highly comparable from both arterioles and venules. Thus, we have developed a single-vessel fMRI platform to map the BOLD, CBV, and CBF from penetrating microvessels throughout the cortex. This high-resolution single-vessel fMRI mapping scheme not only enables the vessel-specific hemodynamic mapping in diseased animal models but also presents a translational potential to map vascular dementia in diseased or injured human brains with ultra-high field fMRI.

**Summary:** We established a high-resolution PC-based single-vessel velocity mapping method using the high field MRI. This PC-based micro-vessel velocity measurement enables the development of the single-vessel CBF-fMRI method. In particular, in contrast to the arteriole-dominated CBV and venule-dominated BOLD responses, the CBF-fMRI shows similar velocity changes in penetrating arterioles and venules in activated brain regions. Thus, we have built a noninvasive single-vessel fMRI mapping scheme for BOLD, CBV, and CBF hemodynamic parameter measurements in animals.

## Introduction

Cerebral blood flow (CBF) is a key readout of neuronal processing and viability in normal and diseased brain states^1^. Changes in CBF may be monitored directly within individual blood vessels through the use of optical-based particle tracking techniques^2^. A variety of imaging methods have been developed to measure CBF across multiple spatial scales from capillary beds to the vascular network in animal brains, including multi-photon microscopy^3^, near-infrared spectroscopy (NIRS)^4^, optical coherence tomography^5^, optoacoustic imaging^6^, or laser doppler and speckle imaging^7, 8^. In particular, the doppler-based functional ultrasound imaging method provides a unique advantage to detect the CBF in the brain with a high spatiotemporal resolution, which can be readily applied for awake animal imaging^9-11^. However, these methods share a common barrier that the spectrum-specific signal transmission cannot effectively pass the skull of animals without significant loss of the signal-to-noise ratio (SNR). Typically, a craniotomy or procedure to thin the skull is needed to detect the hemodynamic signal^2^. While current techniques support transcranial imaging into the superficial layers of the cortex, only functional MRI (fMRI) provides a noninvasive approach for measuring hemodynamic signals throughout the brain.

Changes in CBF may be detected by fMRI based on arterial spin labeling (ASL), in which water protons in a major upstream vessel are spin-polarized with an external field^12-14^. Two other fMRI-based techniques provide indirect information about changes in CBF. Blood oxygenation level-dependent (BOLD) fMRI is used to determine changes in the ratio of deoxy- to oxyhemoglobin in the blood and is a measure of changes in brain metabolism^12, 15, 16^. Cerebral blood volume (CBV) fMRI is used to measure changes in blood volume, i.e., essentially changes the diameter of arterioles, based on the use of exogenous or endogenous contrast agents to differentiate blood from brain tissue^12, 17^.

Phase-contrast (PC) MRI relies on gradient-oriented dephasing of magnetized protons to map the velocity, i.e., direction and speed, of blood flow^18, 19^. The ASL-based CBF fMRI technique detects local changes in the flow of blood through brain tissue but does not show orientation-specific information related to the alignment of vessels^20^. Past works with 7 T MR scanning showed that PC-MRI can be used to measure flow in the perforating arteries through the while matter or the lenticulostriate arteries in the basal ganglia of human brains^21-24^. However, the SNR was insufficient in these prior studies to map changes in flow, and thus changes in CBF.

Here, we report on a PC-MRI method to detect the vessel-specific changes in blood velocity in single trials. Compared with past implementations of PC-MRI^21, 25-28^, we have implemented a small surface radio frequency (RF) coil with the high field MRI, i.e., 14.1 T for improved SNR. This further allows us to map the BOLD- and CBV-fMRI from individual penetrating venules and arterioles, which span 20 to 70 µm diameter, with high spatial resolution^29-31^.

## Results

### Phantom validation of high-resolution PC-based flow velocity measurement

For calibration, we constructed an *in vitro* capillary tubing circulatory system to mimic penetrating vessels, with flow rates from 1 to 10 mm/s (**Figure 1A**). A 2D PC-MRI slice is aligned perpendicular to the capillary tubing (**Figure 1A, B)** and provides a voxel-specific measurement of the flow velocity through two tubes with the upward flow (positive sign, bright dots in **Figure 1B**) and two tubes with the downward flow (negative sign, dark dots in **Figure 1B**), as well as a control tube. We observe a monotonic and near-linear relation between the velocity measured by PC-MRI and the true velocity: V_meas_ = (0.67 ± 0.01) _Vpump_ + (0.02 ± 0.11) mm/s at echo time (TE) = 5.0 ms (**Figure 1C**). The small offset could be caused by eddy current effects and other gradient-related scaling errors of the PC-MRI sequence^32-34^. We further observe that the measured velocities are relatively insensitive to the value of TE (**Figure 1C**).

**Figure 1.**
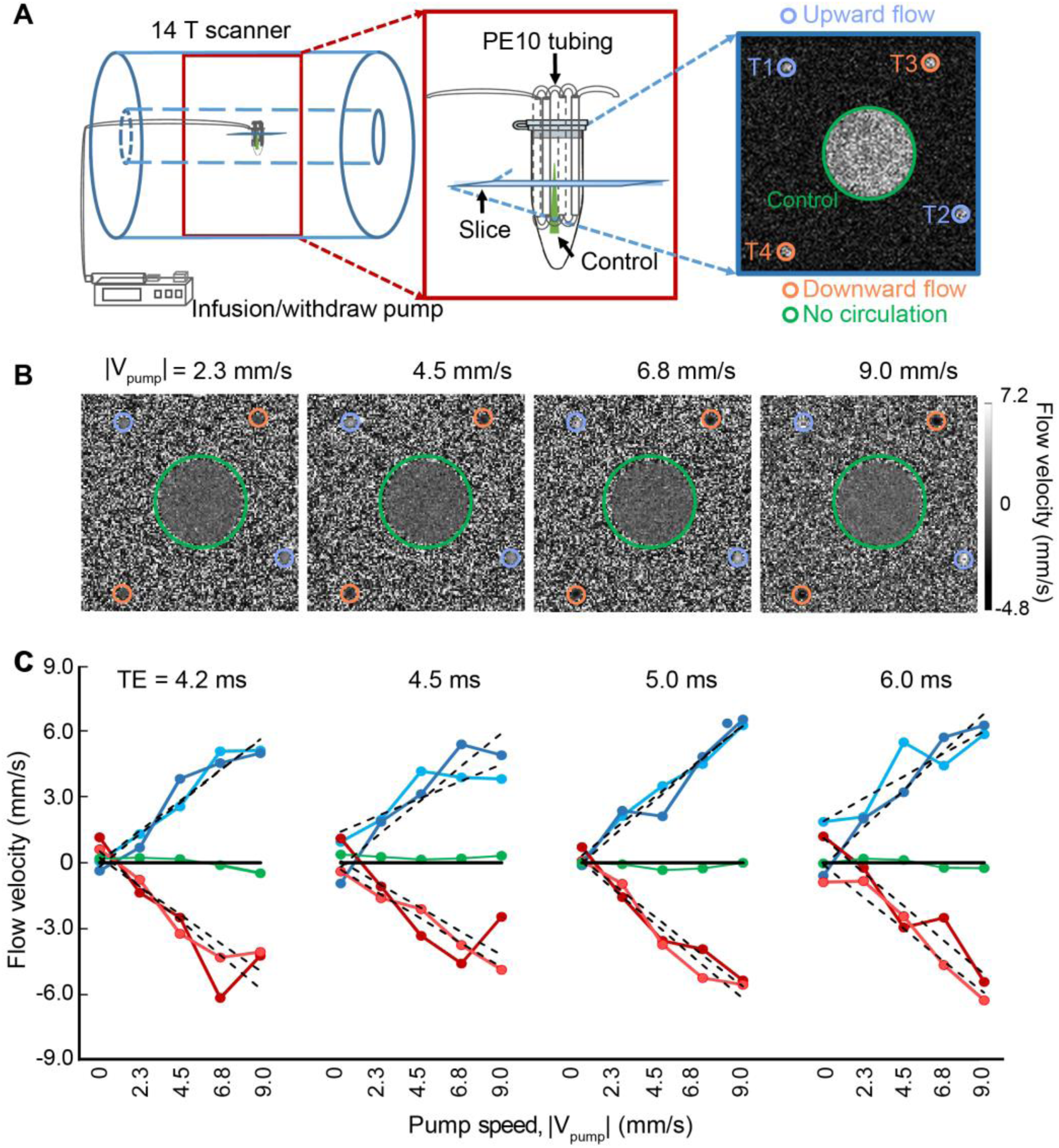
*In vitro* flow velocity measurements with phase-contrast (PC)-MRI. **A**. Schematic drawing of the phantom experimental flow chamber in the 14.1 T scanner. An expanded image (red box) shows the circulatory system constructed of capillary tubes. A representative fast low angle shot (FLASH) MRI image (blue box), 500 µm in thickness, shows the capillary positions. regions of interest (ROIs) T1 and T2, contoured in purple, indicate the upward flow. ROIs T3 and T4, in orange contour, indicate the downward flow. The green contour indicates the stagnant fluid. **B**. Representative images with different flow velocity in the capillaries T1 to T4 in panel A. (echo time) TE = 5.0 ms for all panels. **C**. The plot of flow velocity estimates from the five ROIs with different TEs, as marked, and different pump rates, as indicated and marked in panel B. The dotted lines correspond to a linear fitting for velocity measurements of different ROIs.

We implemented the high-resolution PC-MRI for *in vivo* measurement of blood flow from individual penetrating arterioles and venules through the infragranular cortex, i.e., layer V, of the anesthetized rats with 14.1 T MR scanning. To improve the SNR of PC-MRI images as well as multi-gradient echo (MGE) images used for arteriole-venule (A-V) mapping^29, 31^, a surface RF transceiver coil with 6 mm diameter was constructed and attached to the rat skull (**Supplemental Figure 1**). This was essential for the high-resolution mapping with a fast sampling rate of the single-vessel flow velocity over a complete hemisphere of the rat brain (**Figure 2A and Supplemental Figure 1E**).

**Figure 2.**
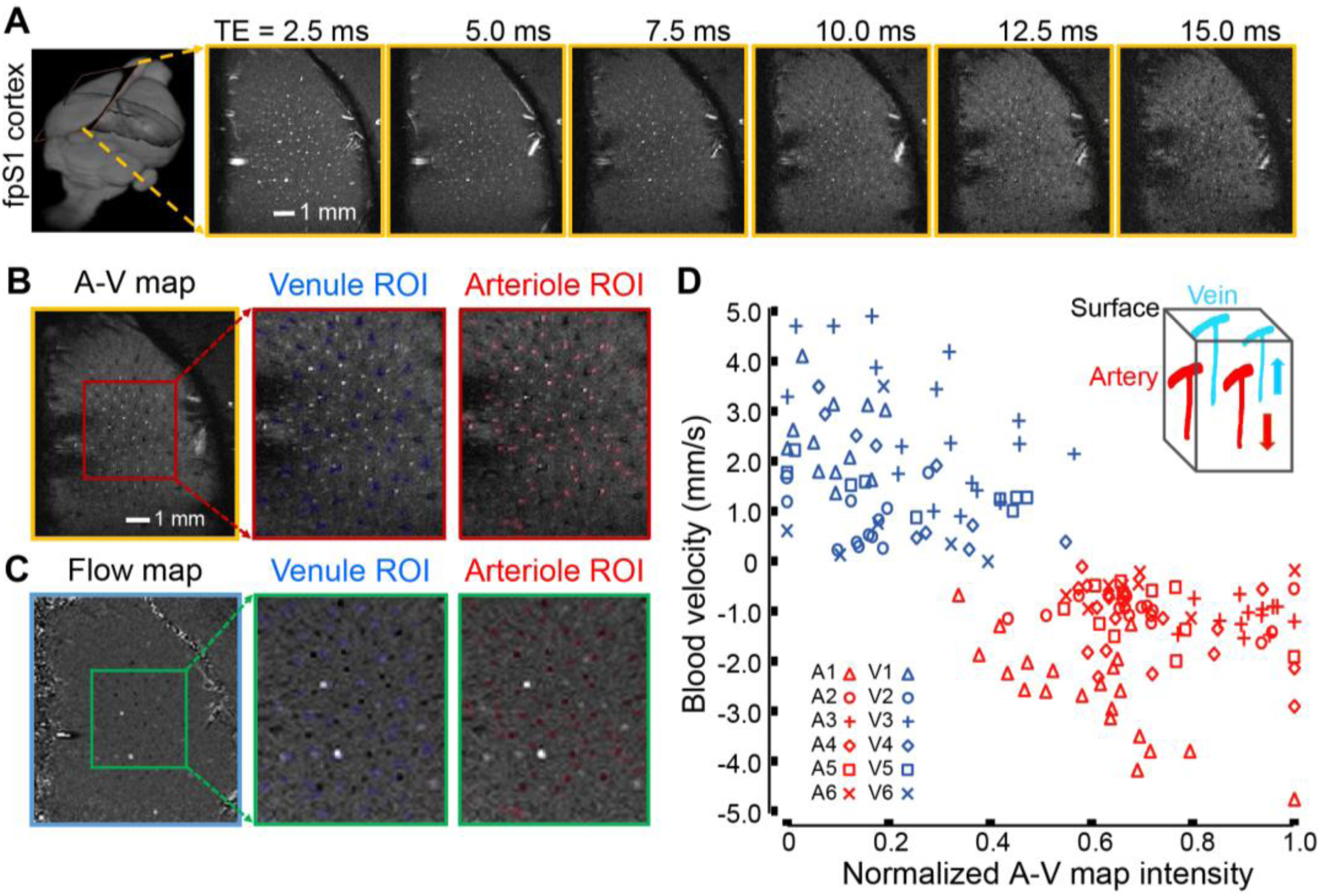
Single-vessel flow velocity measurement. **A**. Representative 2D multi-gradient echo (MGE) slices (yellow boxes) from a deep layer of the primary forepaw somatosensory cortex (first frame) at different TEs, as indicated. **B**. The 2D arteriole-venule (A-V) map (yellow box) derived from the images with different TEs in panel A, arterioles and venules appear as bright and dark voxels, respectively. The expanded views (red boxes) show individual venules, i.e., black voxels marked in blue, and arterioles, i.e., white voxels marked in red. **C**. The vectorized flow velocity map (blue box) from the same 2D MGE slice in panel B. The expanded views (green boxes) show the individual venules, i.e., white dots with positive velocity, and arterioles, i.e., black dots with negative velocity. Note that 2 bright dots are caused by the “over-flowed” velocity beyond the maximal velocity, i.e., the velocity encoding (Venc) parameter, defined in the PC-MRI sequence, which could be not correctly estimated. **D**. Scatter plot of the flow velocities from individual arterioles and venules as the function of the normalized signal intensities of each vessel in the A-V map of panel B, data from 6 rats as indicated. Insert shows the blood flow direction of arterioles and venules in the forepaw somatosensory cortical region.

### *In vivo* PC-based flow velocity mapping of penetrating microvessels

We first acquired the single-vessel A-V map by aligning a 500 µm thick 2D MRI slice perpendicular to the penetrating vessels through layer V of one hemisphere (**Figure 2A, B**). We designed the pulse sequence for PC-MRI to achieve the same slice geometry of the A-V map so that the CBF deduced from PC-MRI signals could be overlaid with individual penetrating arterioles and venules in the single-vessel flow velocity map (**Figure 2A, C**). The arteriole blood flows down into the cortex while the venule blood flows upward, which determines the sign of the flow velocity. Vessel-specific velocities were plotted as a function of the normalized signal intensity in the A-V map and corroborated our ability to determine flow velocity specific to arterioles and venules (**Figure 2C, D**). The measured flow velocities range from 1 to 10 mm/s, as previously measured with optical methods^35^. To probe the reliability of the single-vessel MR-based flow velocity method, we compared the velocities detected by PC-MRI methods with different TEs and flip angles (FAs) and observed comparable results across a range of parameters (**Supplemental Figure 2**). All told, these data demonstrate the feasibility of *in vivo* single-vessel blood velocity mapping with the PC-based MRI method.

### PC-based CBF-fMRI from individual arterioles and venules

We contrasted the complementary capabilities of PC-based CBF-fMRI against the signals observed with the balanced steady-state free precession (bSSFP)-based single-vessel BOLD- and CBV-fMRI mapping method^29^ (**Figure 3**). We first created an A-V map through the deep layers of the forepaw region of the primary somatosensory cortex (**Figure 3Ai**), followed by 2D-bSSFP to detect stimulus-induced changes in the single-vessel BOLD-fMRI signal (**Figure 3Aii**). We next performed single-vessel PC-MRI flow velocity measurements with 100 x100 µm^2^ in-plane resolution, a sampling rate of 4 s repetition time (TR) per image, and the same geometry as the 2D-bSSFP method to measure baseline flow in penetrating arterioles and venules (**Figure 3Aiii**). Changes in CBF upon stimulation overlapped with individual penetrating vessels in the A-V map (**Figure 3Aiv**). Lastly, we performed 2D-bSSFP for single-vessel CBV-fMRI mapping by intravenous injection of iron particles into the blood in the same rats (**Figure 3Av**). The BOLD-fMRI signal is primarily detected from individual penetrating venules while the CBV-weighted signal is mainly located at the individual penetrating arterioles (dark dots in **Figure 3Aii** with bright dots in **Figure 3Av**). In contrast, the CBF-fMRI signal is observed in both penetrating arterioles and venules (**Figure 3Aiv**).

**Figure 3.**
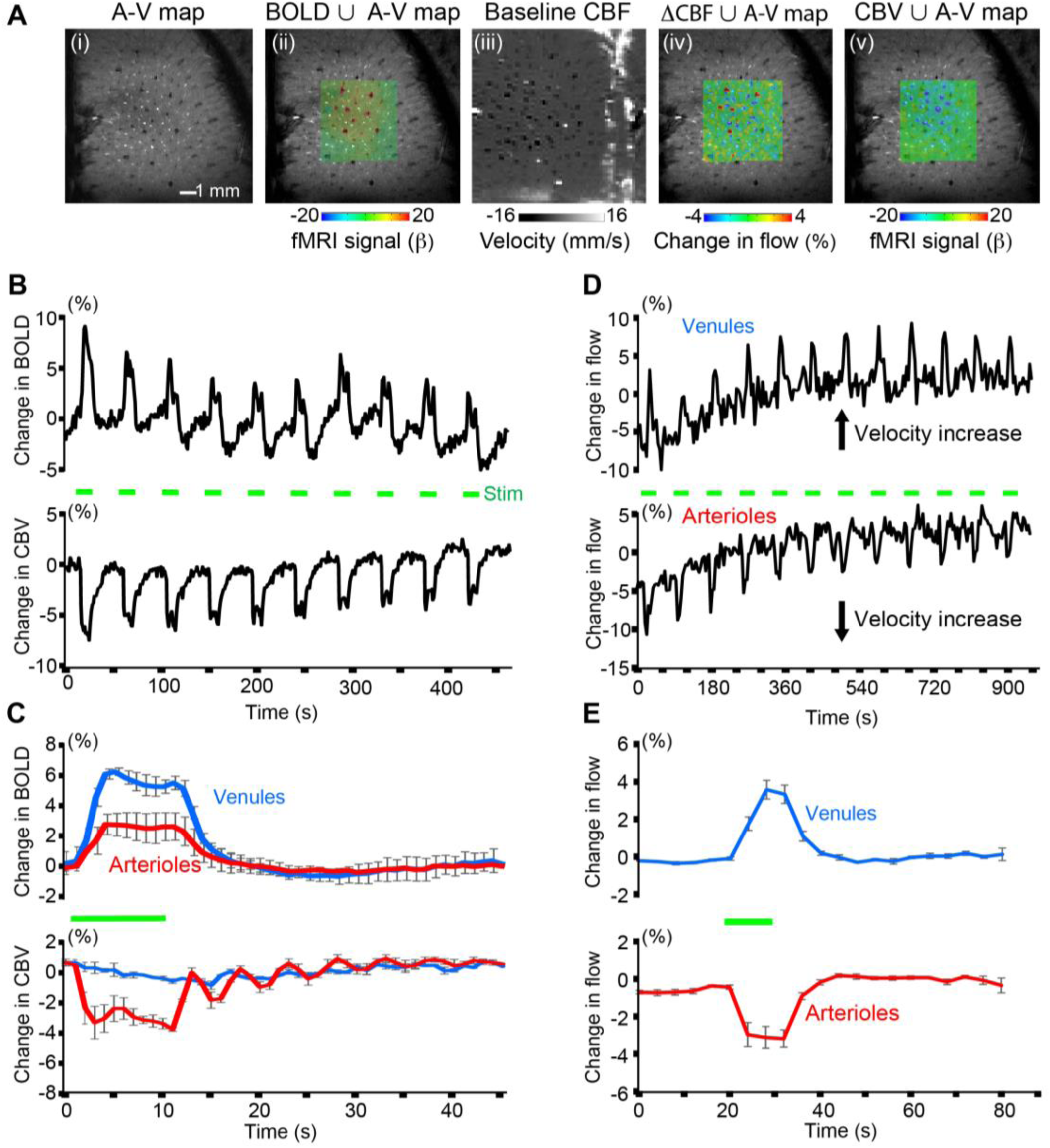
Maps of task-related hemodynamic signals with BOLD, CBV, and CBF-fMRI. **A**. Different MRI measurement strategies on the same 2D slice. From left to right: (i) The single-vessel A-V map acquired with the MGE method; (ii) The evoked balanced steady-state free precession (bSSFP)-based BOLD-fMRI signal, within the green subregion, on top of the A-V map; (iii) the PC-MRI map of baseline CBF; (iv) the change in CBF on top of the A-V map with an increased flow velocity corresponding to brighter voxels for venules and darker voxels for arterioles; and (v) the evoked bSSFP-based CBV-fMRI signal on top of the A-V map. **B**. The time courses of the evoked bSSFP-BOLD and CBV-fMRI with the block-design paradigm from the venules and arterioles shown in panel A. Forepaw stimulation pulse of 330 μs in width and 1 mA in amplitude delivered at 3 Hz for 10 s. **C**. The averaged time courses of the fractional change for the evoked BOLD and CBV signals from the venule and arteriole ROIs from different rats (mean ± SEM, the green bar shows stimulation duration). The peak BOLD values of venule are significantly higher than those of arteriole (5 rats, p = 0.009), while the peak CBV values of arteriole are significantly higher than those of venule (3 rats, p = 0.028). **D**. The time courses of the evoked CBF changes from the arteriole and venule ROIs show increased velocity from both arterioles and venules with the block–design, 10 s duration stimulation paradigm. **E**. The averaged time courses of the evoked CBF changes show the velocity increase from both arteriole and venule ROIs with the block–design stimulation paradigm from 4 rats (mean ± SEM, the green bar shows stimulation duration).

The stimulus-evoked responses of all three fMRI signals were studied with an on/off block design (**Figures 3B-E**). Group analysis shows that the positive BOLD signal from venule voxels is significantly higher than the arteriole-specific BOLD signal (**Figure 3C**). In contrast, the arteriole dilation leads to an earlier CBV-weighted negative fMRI signal, which is much stronger and faster than the signal from passive venule-dilation (**Figure 3C**), as expected^36, 37^. Group analysis shows the similar temporal dynamics of CBF changes appearing as a ramp in both arterioles and venules (**Figure 3E**). The CBF-fMRI signal appears as the integral of the stimulus, i.e., a triangular ramp (**Figure 3E**), compared to the saturation-like BOLD and CBV responses (**Figure 3C**). The voxel-wise hemodynamic changes of BOLD, CBV, and CBF are illustrated in **Supplementary Figure 3** and **movie 1**.

## Discussion

Despite the existing tools developed for CBF measurement in both animal and human brains, it remains challenging to detect the flow dynamics of intracortical micro-vessels non-invasively. Here, we not only optimized the PC-MRI to map the vectorized single-vessel flow velocity of penetrating arterioles and venules but also developed the single-vessel CBF-fMRI based on the direct flow velocity measurement in rat brains. By combining with previously established single-vessel BOLD- and CBV-fMRI methods, the PC-based single-vessel CBF-fMRI method complements the scheme to map the vessel-specific hemodynamic responses with high-resolution fMRI.

In contrast to the conventional ASL methods, PC-based MRI mapping allows arterioles and venules to be distinguished for simultaneous velocity measurements through a 2D plane. Also, ASL has less vascular specificity because water exchange through the blood-brain barrier of capillary beds increases the weighting of the ASL-based flow signal for parenchyma voxels^38, 39^. Furthermore, there is significant variability in the transit time to flow from arterioles to venules through the capillary bed^40^, which complicates the distinction of arterioles and venules by simple ASL-based CBF mapping. We detected the velocity from penetrating microvessels in the deep cortical layers with PC-MRI, showing velocity values from 1 to 5 mm/s (Fig 2). Single-digit velocity (mm/s) of red blood cells from cortical surface microvessels has been detected in the anesthetized rats using two-photon microscopy^2, 35^. It is noteworthy that the PC-based vessel velocity measurement is based on measuring water protons in blood but not limited to the flow of red blood cells. Still, the PC-based velocity from microvessels matches well with the previous optical measurement. We conclude that high-resolution PC-MRI is ideal for noninvasive single-vessel CBF-fMRI mapping.

A remaining complication with PC-MRI mapping is the presence of small offsets in velocity from the phantom capillary tubing with circulating flow under different conditions. The phase-dependent velocity encoding depends on the quality of the magnetic field gradients, and mismatched eddy currents of multiple gradients with opposite polarities, as well as the nonlinear and distorted gradient fields, could contribute to distortions in gradients^32-34^. In particular, the high-resolution PC-MRI method is a high-duty cycle sequence and slight heating of the gradient coil during scanning may alter the gradient performance, consistent with the baseline-drift of the CBF-fMRI signal in the first 5 minutes of scanning (**Figure 3D**). Nevertheless, it should be noted that the percentage velocity changes from individual arterioles and venules can be readily detected with the PC-based CBF-fMRI measurement regardless of the gradient-heating related baseline drift. Another factor that contributes to the phase-dependent velocity error originated from the limited spatial resolution of the PC-MRI images compared to the diameters of small vessels^27^, although corrections are possible^21, 28^. Despite the potential partial volume contribution to the single-vessel BOLD, CBV, and CBF-fMRI, the vessel-specific mapping scheme presents a translational potential to identify vascular dementia in diseased or injured brains with ultra-high field fMRI.

## ACKNOWLEDGMENTS

We thank Dr. E. Weiler, Ms. H. Schulz, and Ms. S. Fischer for animal/lab maintenance and support, Dr. K. Buckenmaier and Dr. N. Avdievitch for technical support, and the Analysis of Functional NeuroImages (AFNI) team for their software support. This research was supported by NIH BRAIN grants R01 1NS113278 and R01 MH111438, NIH NINDS grant R35 NS097265, NIH instrument grant S10 RR023009 to the Massachusetts General Hospital/Harvard-MIT Program in Health Sciences and Technology Martinos Center, Deutsche Forschungsgemeinschaft (DFG, Germany Research Foundation) grant YU 215/3-1, SCHE 658/15, SCHE 658/12, the Bundesministerium fuer bildung und forschung (BMBF, Federal Ministry of Education and Research) grant 01GQ1702, and the Chinese Scholarship Council for the doctoral support of X. Chen.

## AUTHOR CONTRIBUTIONS

X.Y. and D.K. initiated the concept; X.Y. designed the research; X.C. and X.Y. performed animal experiments; X.C., Y.J. performed data analysis; P.R., K.S., S.C. provided technical support; X.Y., D.K., X.C., Y.J. wrote the paper.

## Methods

### Design of a phantom capillary tubing flow system

In order to validate the PC-MRI sequence, a plastic circulatory flow phantom composed of the capillary tubing (PE-10, Instech Laboratories, inner diameter 210 µm) was constructed to mimic the geometries of cortical blood vessels (**Figure 1A**). The capillary tube was connected to a programmable syringe infusion/withdraw pump (Pump Elite 11, Harvard Apparatus) with an infusion rate of 0.25, 0.5, 1.0, 1.5, 2.0 ml/h, which were transferred to the flow velocity of the capillary tubing as shown in **Figure 1**. The flowing medium is a manganese solution (50 mM MnCl_2_, Sigma-Aldrich). The phantom tube was cast with Fomblin (Sigma-Aldrich) to avoid the potential air interface artifacts.

### Animal preparation for fMRI

All surgical and experimental procedures were approved by the local authorities (Regierungspraesidium, Tübingen Referat 35, Veterinärwesen, Leiter Dr. Maas) and were in full compliance with the guidelines of the European Community (EUVD 86/609/EEC) for the care and use of laboratory animals. The experimental animals were Sprague-Dawley male rats, ∼ 250 g, provided by the Charles River Laboratories in Germany. Fifteen rats were used in all experiments (the evoked bSSFP-BOLD/CBV and PC-MRI signals were acquired from five of these fifteen rats).

Detailed descriptions of the surgery are given in previous publications^29, 30^. Briefly, rats were first anesthetized with isoflurane (5% induction, 1∼2% maintenance), each rat was orally intubated with a mechanical ventilator (SAR-830, CWE). The femoral artery and vein were catheterized with plastic catheters (PE-50, Instech Laboratories) to monitor the arterial blood gas, administrate drugs, and constantly measure the blood pressure. After catheterization, rats were secured in a stereotaxic apparatus, a custom-made RF coil was fixed above the skull with cyanoacrylate glue (454, Loctite). After surgery, isoflurane was switched off and a bolus of α-chloralose (80 mg/kg, Sigma-Aldrich) was intravenously injected. A mixture of α-chloralose (26.5 mg/kg/h) and the muscle relaxant (pancuronium bromide, 2 mg/kg/h) was continuously infused to maintain the anesthesia and minimize the motion artifacts. Throughout the whole experiment, the rectal temperature of rats was maintained at 37°C by using a feedback heating system. All relevant physiological parameters were constantly monitored during imaging, including heart rate, rectal temperature, arterial blood pressure, the pressure of the tidal ventilation, and end-tidal CO_2_. Arterial blood gases were checked to guide the physiological status adjustments by changing the respiratory volume or administering sodium bicarbonate (NaBic 8.4 %, Braun) to maintain normal pH levels. Dextran-coated iron oxide particles (15 ∼ 20 mg of Fe/kg, BioPAL, MA) were intravenously injected for CBV-weighted signal acquisition.

### fMRI setup

All images were acquired with a 14.1 T, 26 cm horizontal bore magnet (Magnex Scientific) interfaced through the Bruker Advance III console (Bruker Corporation). The scanner is equipped with a 12 cm magnet gradient set capable of providing a strength of 100 G/cm and a 150 μs rise time (Resonance Research Inc.). A custom-made transceiver coil with an internal diameter of 6 mm was used for fMRI images acquisition. For the electrical stimulation, two custom-made needle electrodes were placed on the forepaw area of the rats to deliver the electrical pulse sequences (330 μs duration at 1.0 ∼ 2.0 mA. The pulses repeated at 3 Hz for 10 s) by using a stimulus isolator (A365, WPI). The stimulation duration and frequency were triggered directly through the MRI scanner which controlled by Master-9 A.M.P.I system (Jerusalem, Israel). The triggering pulses from the MRI scanner were also recorded by the Biopac system (MP150, Biopac Systems, USA).

### Single-vessel multi-gradient echo (MGE) imaging

To anatomically map the individual arterioles and venules penetrating the deep cortical layers of the somatosensory cortex, a 2D MGE sequence was applied with the following parameters: TR = 50 ms; TE = 2.5, 5.0, 7.5, 10.0, 12.5 and 15.0 ms; flip angle (FA) = 55°; matrix = 192 × 192; in-plane resolution = 50 × 50 μm^2^; slice thickness = 500 μm. The A-V map was made by averaging the MGE images from the second TE echo to the fifth TE echo. In the A-V map, the arteriole voxels show bright (red marks) due to the in-flow effect and venule voxels show as dark dots (blue marks) because of the fast T_2_* decay (**Figure 2B**).

### Balanced steady-state free precession (bSSFP) BOLD- and CBV-fMRI

The bSSFP sequence was applied to acquire the evoked BOLD signals by using the following parameters: TR = 15.6 ms; TE = 7.8 ms; flip angle = 15°; matrix = 96 × 96; FOV = 9.6 × 9.6 mm; in-plane resolution = 100 × 100 μm^2^; slice thickness =500 μm. For the bSSFP CBV-fMRI, the parameters were adjusted with TR = 10.4 ms and TE = 5.2 ms. The total TR to acquire each image is 1 s. To reach the steady-state, 300 dummy scans were used, followed by 25 pre-stimulation scans, one scan during stimulation, and 44 post-stimulation scans with 10 epochs for each trial. The fMRI stimuli block design of each trial consisted of 10 s stimulation and 35 s inter-stimulus interval. The total acquisition duration of each trial was 7 min 55 s. CBV-weighted fMRI signals were acquired after intravenous injection of dextran-coated iron oxide particles (15 ∼ 20 mg of Fe/kg, BioPAL, MA).

### Phase Contrast (PC)-MRI

To measure the flow velocity of individual arterioles and venules, the PC-MRI sequence was applied with the following parameters. For the *in vitro* phantom measurement: TR = 15.6 ms; TE = 4.2, 4.5, 5.0, 6.0 ms; flip angle = 25°; FOV = 6.4 × 6.4 mm; matrix = 128 × 128; in-plane resolution = 50 × 50 μm^2^; slice thickness = 500 μm; maximum velocity (Venc) = 1.56 or 0.66 cm/s (based on the flow values); number of averages = 172. The total acquisition time was 11 min 28 s. For the *in vivo* measurements: TR = 15.6 ms; TE = 5 ms; flip angle = 30°; FOV = 6.4 × 6.4 mm; matrix = 64 × 64; in-plane resolution = 100 × 100 μm^2^; slice thickness = 500 μm. A total TR for each image is 4 s. The total acquisition duration of each trial was 16 min. To measure the blood flow velocity, bipolar flow encoding gradients were applied along the slice encoding direction. The slice position was anatomically identical with the slice position of the MGE imaging.

### Data analysis and statistics

All data preprocessing and analysis were performed by using the software package, Analysis of Functional NeuroImages (AFNI) (NIH, Bethesda). All relevant fMRI analysis source codes can be downloaded from https://www.afni.nimh.nih.gov/afni/.

### Definition of the individual vessels

The individual arteriole/venule voxels were defined by the signal intensity of the A-V map^31^. The arterioles are determined if the voxel intensities are higher than the mean signal intensities plus two times the standard deviation of the local area in a 5 × 5 kernel. The venules are determined if the voxel intensities are lower than the mean signal intensities minus two times the standard deviation of the local area^29-31^. The locations of individual arteriole/venule voxels defined in A-V map were used to extract the time courses of BOLD/CBV-fMRI for individual vessels.

### BOLD/CBV-fMRI and PC-MRI data analysis

To register the evoked bSSFP-fMRI images and evoked PC-MRI images with the 2D anatomical A-V map, the tag-based registration method was applied. Twelve to fifteen tags were selected from the averaged bSSFP-fMRI images or the averaged PC-fMRI images to register those selected from A-V map. We used a 3dLocalstat AFNI function to normalize the signal intensity of the single-vessel maps. This process allowed us to plot the PC-based velocity values of individual vessel voxels to the normalized signal intensity of A-V maps. For the evoked signals, the bSSFP-fMRI images and PC-MRI images were normalized by scaling the baseline to 100. Multiple trials of block-design time courses were averaged for each animal. No additional smoothing step was applied. The β-value was calculated to measure the amplitude of the fMRI responses at each TR. The voxel-wise β-map was illustrated with the spatial pattern of the fMRI responses corresponding to the different time points after the stimulus onset. After registration (tag-based registration) and region of interest extraction (3dLocalstat function, mask shown in **Figure 2B**), we extracted the PC-based flow velocity values from individual vessel voxels, which were identified based on the algorithm as described in the previous section.

The hemodynamic response function is based on the “block function” of 3dDeconvolve module developed in AFNI. The HRF model is defined as follows:

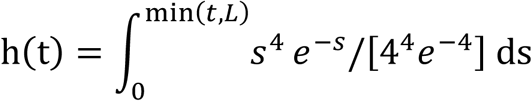

Gamma variate function = s^4^e^-s^ /4^4^e^-4^. L was the duration of the response. BLOCK (L, 1) is a convolution of a square wave of duration L, makes a peak amplitude of block response = 1.

For the group analysis, Student’s t-test was performed, error bars are displayed as the means ± SEM. The p values < 0.05 were considered statistically significant. The sample size of animal experiments is not previously estimated. No blinding and randomization design was needed in this work.

## Supporting information

**Figure S1.**
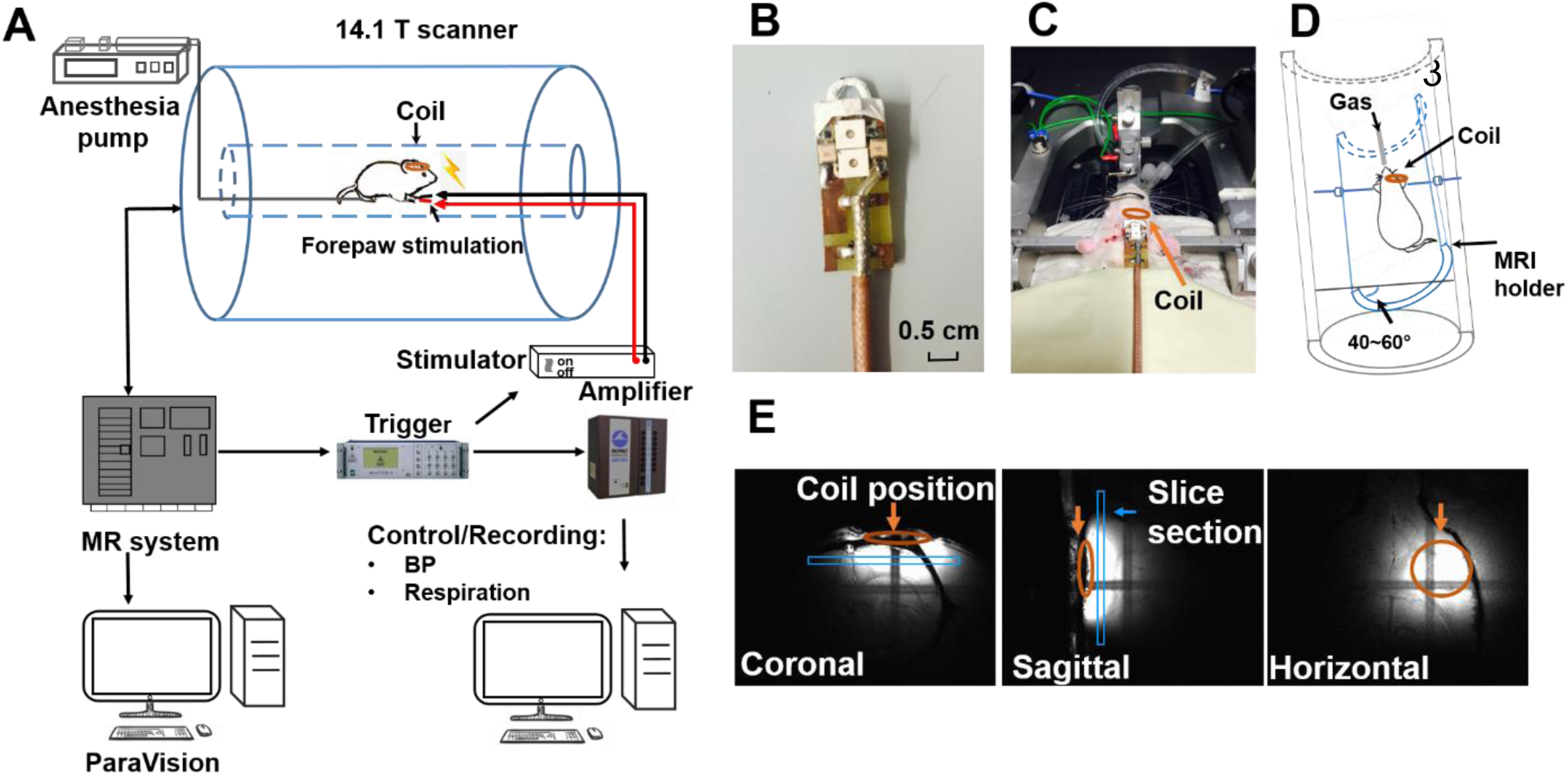
The preparation of *in vivo* experiment for the PC-MRI in 14.1 T. **A**. The flow chart of the *in vivo* experiment in the 14.1 T scanner. **B**. Photograph of the custom-made transceiver surface RF coil. **C**. Photograph of the coil position: the coil is glued to the rat skull. **D**. The schematic drawing of the rat position inside the MRI holder. **E**. Representative images from different views of the FLASH MRI show the ideal coil position.

**Figure S2.**
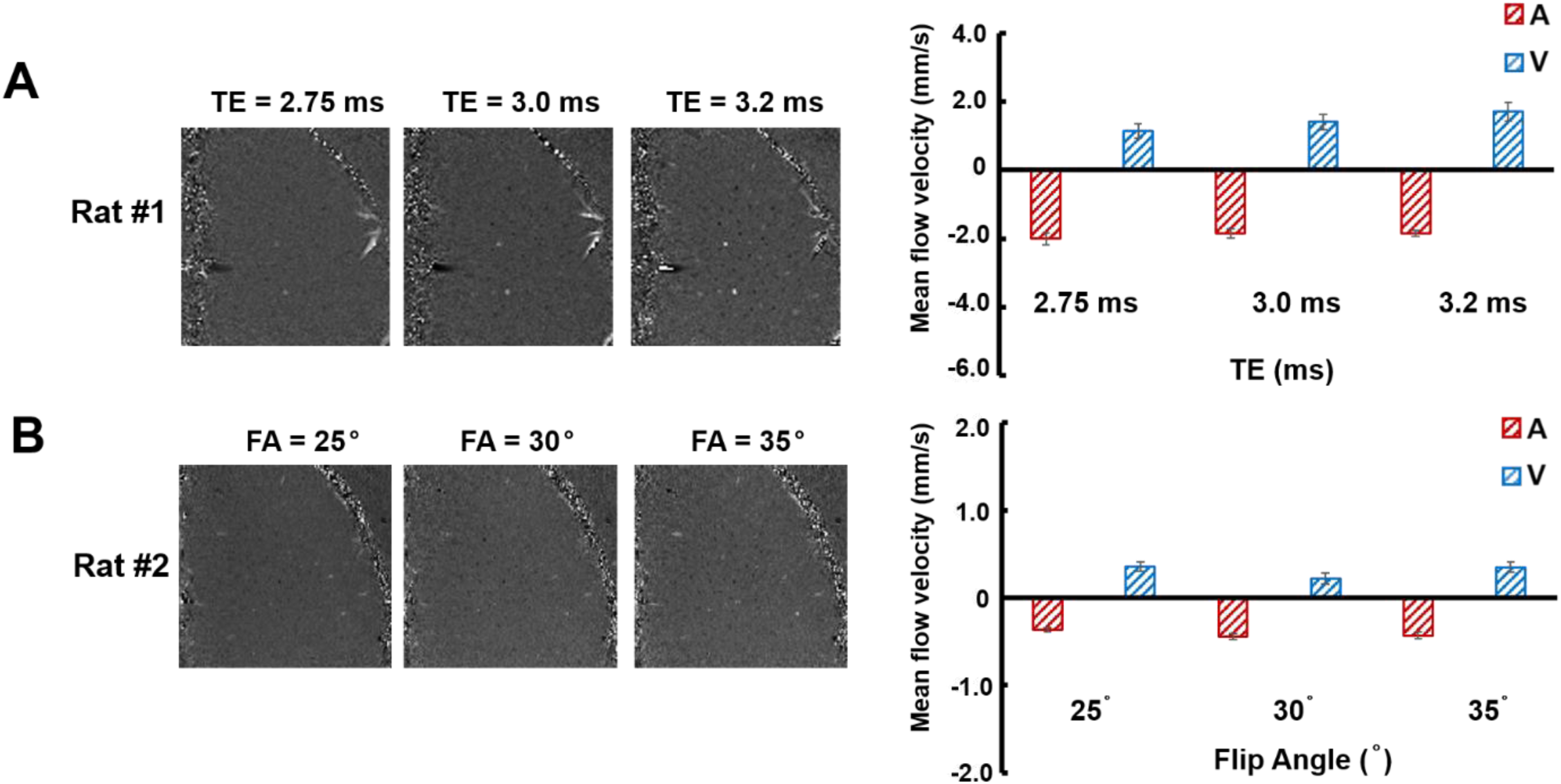
Phase images from the different representative rats with different TEs and flip angles. **A**. Phase images from a representative rat with different TEs, i.e., 2.75, 3.0, and 3.2 ms. The right panel shows the mean blood flow velocity (mean ± SEM) from left images with N_Arteriole_ = 48 and N_Venule_ = 22. **B**. Phase images from a representative rat with different flip angles, i.e., 25°, 30°, and 35°. The right panel shows the mean blood flow velocity from left images with N_Arteriole_ = 38 and N_Venule_ = 14.

**Figure S3.**
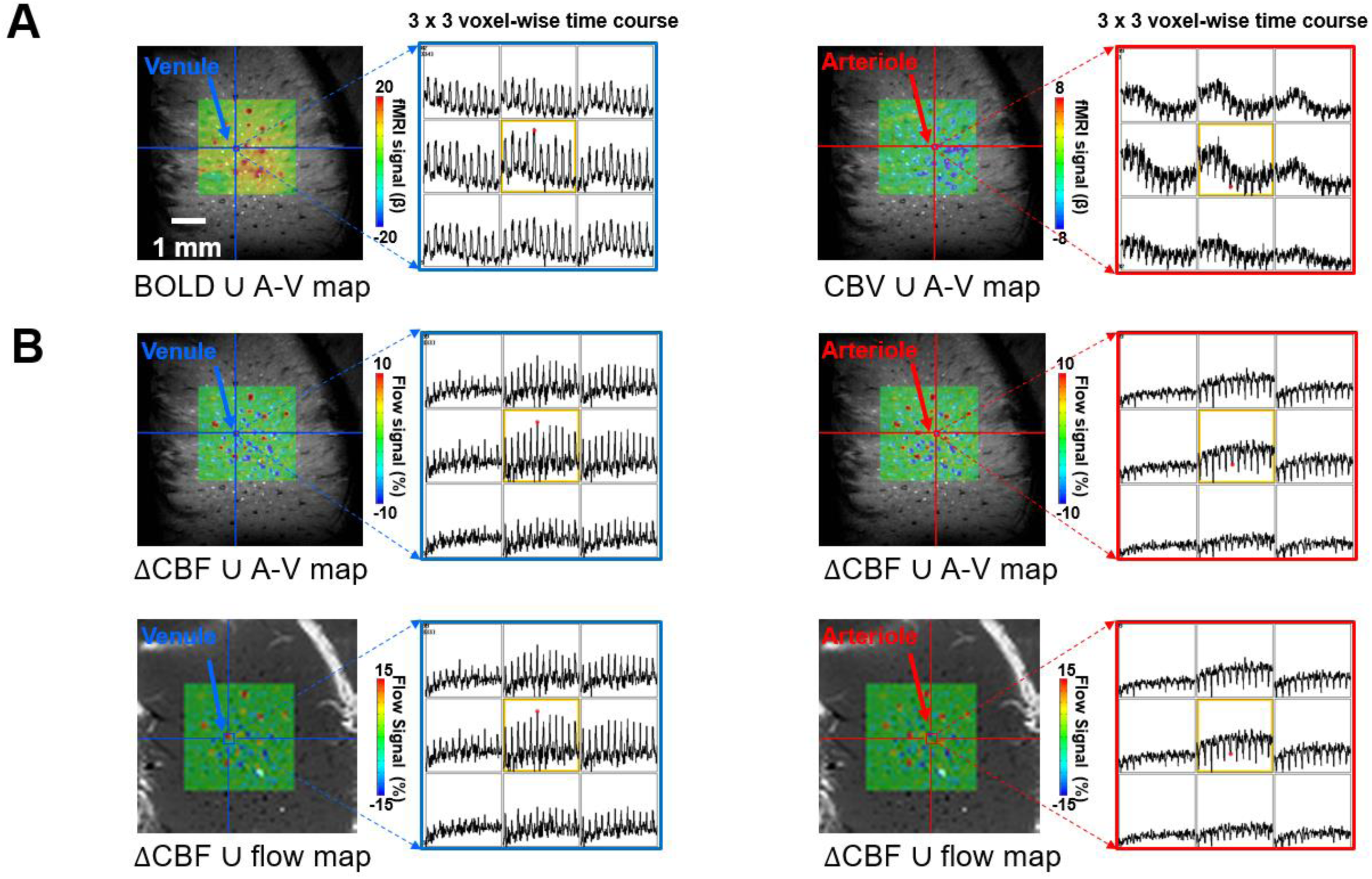
The bSSFP-based single-vessel BOLD/CBV-fMRI and the PC-MRI based single-vessel dynamic flow measurement from a representative rat. **A**. The evoked bSSFP-based BOLD-(left) and CBV-(right) fMRI maps overlaid on the same A-V map from a representative rat, with the voxel-wise time courses from the ROIs of individual venule (blue arrow) and arteriole (red arrow) (10 s on and 35 s off for 10 epochs plotted in a 3 × 3 matrix). **B**. The evoked CBF maps overlaid on both A-V map (upper panel) and PC-based flow map (lower panel) from the same representative rat. The voxel-wise time courses of CBF changes from the same ROIs of individual venule (blue arrow) and arteriole (red arrow) (10 s on and 50 s off for 12 epochs plotted in a 3 × 3 matrix).

## Supplementary Movie Legends

**Movie 1. The bSSFP-based single-vessel BOLD/CBV-fMRI and the PC-MRI based single-vessel dynamic flow measurement in the rat cortex**.

The upper panel shows the evoked bSSFP-based BOLD-(left) and CBV-(right) fMRI maps overlaid on the 2D A-V map with 50 × 50 µm^2^ in-plane and the time course from a single voxel located at a representative venule (blue arrow) and arteriole (red arrow) (10 s on and 35 s off for a total of 45 s time window with TR = 1 s). The middle panel shows the evoked CBF maps overlaid on the same 2D A-V map from the same individual venule (left) and arteriole (right), of which the CBF-fMRI maps were registered to match the 2D A-V map at the 50 × 50 µm^2^ resolution, as well as the time courses of the PC-based flow velocity dynamic changes from same voxels identified as the upper panel (10 s on and 50 s off for a total of 60 s time window with TR = 4 s). The lower panel shows the evoked CBF maps overlaid on the PC-based flow map with 100 × 100 µm^2^ resolution and the time course from the voxels located at the representative venule (left) and arteriole (right). Note that the venule is a bright voxel and arteriole is a dark voxel in the flow map, which is opposite to the A-V map and also the slightly different time course due to the altered spatial resolution.

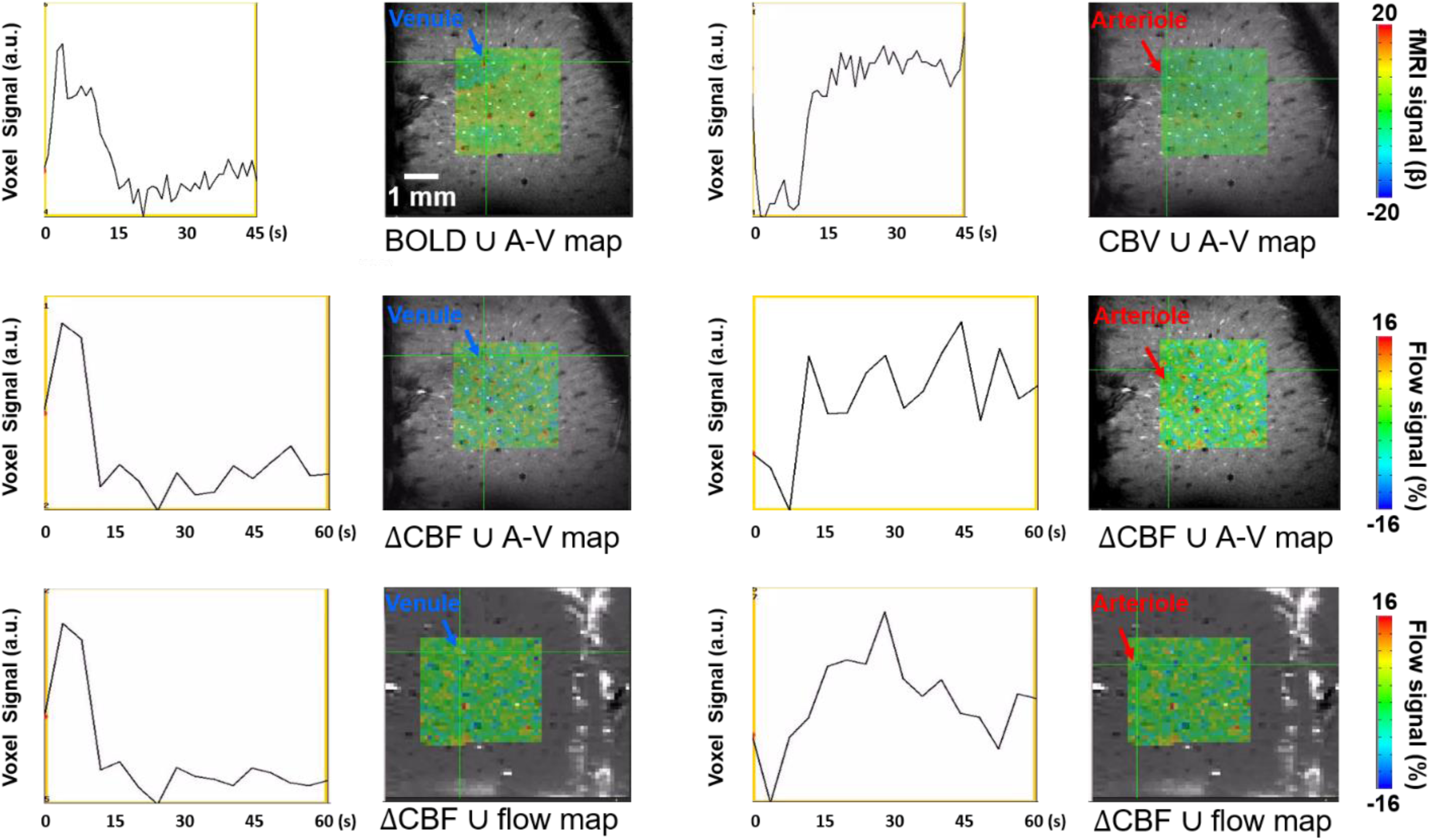

